# Coupling and Decoupling between Brain and Body Oscillations

**DOI:** 10.1101/484097

**Authors:** Elie Rassi, Georg Dorffner, Walter Gruber, Wolfgang Klimesch

## Abstract

Cross frequency coupling is used intensively to study the cross talk between brain oscillations. In this paper we focus on a special type of frequency coupling between brain and body oscillations, which is reflected by the numerical ratio (r) between two frequencies (m and n; n > m). This approach is motivated by theoretical considerations indicating that an integer relationship (r = n/m = integer number) reflects coupling, whereas an irrational relationship (r = n/m = irrational number) reflects decoupling. We analyzed alpha frequency, heart rate, breathing frequency and spindle frequency from data collected by the SIESTA research group. Our results show a 1:4 frequency relationship between heart rate and breathing frequency both during wakefulness and sleep. During wakefulness we expected but did not find an integer relationship between alpha frequency and heart rate or alpha frequency and breathing frequency. During sleep, we observed an irrational relationship between spindle frequency and heart rate as well as spindle frequency and breathing frequency–suggesting decoupling.

## 1. Introduction

Cross frequency coupling is well documented in numerous publications about brain oscillations (for reviews see e.g., Jensen & Colgin, 2007, Canolty & Knight, 2010; Palva & Palva, 2017), body oscillations (specifically heart rate variability; Acharya et al. 2007) as well as brain and body oscillations (see e.g. the ‘network physiology approach’, Bashan et al. 2012, or studies focusing on specific oscillations, such as e.g., gastric waves and alpha, Richter et al. 2017). Physiologically, there are two well described coupling principles that govern brain and body oscillations. Amplitude (envelope) coupling between any frequencies m and n, where the phase of the slower frequency m modulates the envelope of the faster frequency n, and phase coupling between m and n. Amplitude and phase coupling differ with respect to at least the following two properties.

(i)Phase coupling requires a harmonic (integer) relationship, but amplitude coupling works for any m:n frequency ratio. This is a simple but interesting fact, showing that cross frequency phase coupling can be considered a ‘two step’ process: Two oscillations shift their frequencies (m, n) in a way that their ratio is an integer (r = m/n = integer number), which then, in a second step, invites phase coupling. On the other hand, if the ratio produces an irrational number (r = m/n = irrational number), stable phase coupling is disabled (Pletzer et al. 2010; Roopun et al. 2008). As an illustration, let us assume three sine oscillations, one with 10 Hz, another with 20 Hz, and a third with g*10 = 16.18… Hz, where g is an irrational number, termed golden mean (g = 1.618….). Let us further assume that the positive peaks of the three oscillations are exactly aligned to each other at time t(0). Then, the positive peaks of the 10 Hz and 20 Hz oscillations will meet regularly, at any second peak of the faster and at each peak of the slower oscillation. This means that the two harmonically related oscillations are strictly phase coupled (in this example with zero phase lag). In contrast, the positive peaks of the 10 Hz and the g-related 16.18… Hz oscillations will never meet at any time point t(i) in the future (for a mathematical proof and analysis see Pletzer et al., 2010). The g-related oscillations are de-coupled in a way that their positive peaks vary together in an irregular pattern over time. Pletzer et al. (2010) have also shown that this pattern is most irregular for g, meaning that in this sense g is the ‘most irrational’ number.

(ii)Phase coupling operates at the temporal precision of the faster oscillation, but amplitude coupling operates at the temporal precision of the slower oscillation. The reason is that the excitatory time window of the faster oscillation – which can be driven by the phase of the slower oscillation - is smaller than that of the slower oscillation. This property makes cross frequency phase coupling an interesting candidate for cognitive top down control, which is closely associated with widespread and long range brain synchronization (for an empirical analysis see e.g., Siebenhühner et al. 2016). The hypothesis is that slow oscillations (in the delta, theta, alpha and beta frequency range) play an important role for the downstream control of neuronal synchronization in anatomically distributed neural circuits (Fries, 2015; Palva & Palva, 2017) in a way that the phase of a slow oscillation drives the phase of a fast oscillation (particularly in the gamma frequency range).

Based on the specific properties (and assumed functions) of cross frequency phase coupling, Klimesch (2013; 2018) has suggested that the center frequencies of traditional EEG frequency bands are harmonically related and form a binary hierarchy of frequencies (delta = 2.5 Hz, theta = 5 Hz, alpha = 10 Hz, beta = 20 Hz, gamma1 = 40 Hz), in which the neighboring higher frequency always is twice as fast than its slower neighbor. It is also assumed that this harmonic frequency architecture allows optimal brain communication that typically appears during cognitive processing demands (see e.g., Palva & Palva, 2017). Most importantly, Klimesch (2018) has reviewed evidence showing that this binary hierarchy extends ‘down’ to and allows to predict the frequencies of body oscillations. If we go down this binary hierarchy from e.g., delta with 2.5 Hz, we obtain the following slower frequencies: 1.25 Hz, 0.625 Hz, 0.3125 Hz etc. These frequencies can be associated with heart rate (1.25 Hz), the frequency of muscle contraction supporting inhaling and exhaling (0.625 Hz), and breathing frequency (0.3125 Hz).

In this paper we test predictions of the binary hierarchy brain body oscillation theory as suggested by Klimesch (2013; 2018). We ask whether the frequency ratios between brain and body oscillations, such as alpha frequency (AF), heart rate (HR) and breathing frequency (BF) vary arbitrarily between subjects or are numerically related to each other, reflecting a binary hierarchy of frequencies. An additional question we ask is whether the expected frequency architecture between brain and body oscillations changes during sleep. To investigate this latter question, we relate sleep spindle frequency (SF), to HR and BF.

The following hypotheses can be derived from the binary hierarchy theory: For AF : HR, AF : BF and HR : BF, we expect frequency ratios of r = 8, 32, and 4 respectively. Or in other words, AF is three binary steps (i.e., 2^3^ = 8) faster than HR and 5 binary steps (2^5^ = 32) faster than BF, whereas HR is two binary steps faster (i.e., 2^2^ = 4) than BF. Studies on mean HR in adults (during wakefulness) are in line with the predicted r = 8, because AF (with a mean value around 10 Hz) is about 8 times faster than HR with a mean value around 1.25 Hz (i.e., 75 beats per minute ; e.g. Shaffer et al. 2014). BF shows multiple peaks, with a prominent peak at about 0.30 Hz (e.g., Perlitz et al. 2004), which is close to the predicted frequency of about 0.31. Thus, the expected ratio between AF and BF equals 32 and the expected ratio between HR and BF equals 4.

For sleep, we can generate further hypotheses that refer to decoupling between brain and body oscillations. The general reason for this assumption is that during sleep, no task related coordination between body actions (such as movements) and brain actions (comprising a variety of different cognitive tasks) is required. We further assume that this situation is characterized by a lack of body awareness. Research on the heartbeat evoked potential (HEP; an evoked potential, calculated time locked to the R peak; first reported in Schandry et al., 1986), which reflects an interesting aspect of body-brain communication, provides empirical backing for this assumption. During alert wakefulness, the magnitude of the HEP response appears to be associated with interoceptive and sensory awareness (Pollatos & Schandry, 2004; Montoya et al., 1993; Schandry & Weitkunat, 1990; Park et al. 2014; Park & Tallon-Baudry, 2014). These findings have led Lechinger et al. (2015) to hypothesize that the magnitude of the HEP response will decrease in sleep. Indeed, the results showed that HEP amplitudes decreased with sleep depth. This study also showed that during wakefulness HR was significantly correlated with AF, but with SF during sleep. The correlation was strongest during wakefulness and declined with increasing sleep depth, suggesting decoupling of brain oscillations from HR. Based on these findings, we expect an irrational number to be produced by the ratios (r = m/n = irrational number) between SF on the one hand and HR and BF on the other hand. Mean SF has a frequency of about 13 Hz (for a review see e.g., De Gennaro & Ferrara, 2003) and a decoupling with HR can be described by SF = g* HR * x. Solving for x shows a value around 8, if we assume that HR drops to around 1 Hz during sleep. Most interestingly, x = 8, represents a binary multiple (2^3^), which according to the binary hierarchy theory, represents a decoupled frequency that is 3 binary steps faster than HR. Thus, in a situation requiring decoupling instead of coupling, the predicted dominant brain frequency shifts from alpha to the spindle frequency range. For body oscillations, there is no reason to assume decoupling during sleep, because body functions must be coordinated during wakefulness and sleep as well. Thus, for sleep, we still expect a 1:4 ratio for HR : BF. If the ratio between HR and BF remains constant and does not change during sleep while SF shifts towards the predicted irrational ratio (of g*HR*8 = 12.94) this also means that the ratio between SF : BF becomes irrational with a predicted value of g*HR*32 = 51.78.

In the present study, we have analyzed AF, HR, BF and SF from data which were collected by the SIESTA project (Klösch et al. 2001). In summarizing our hypotheses, we expect binary multiple frequency ratios for AF : HR, AF : BF and HR : BF during wakefulness, irrational ratios for SF : HR and SF : BF during sleep, but a constant binary multiple ratio for HR : BF.

## 2. Methods

### Subjects

We used recordings and demographic data from the SIESTA project (Klösch et al. 2001), which was established to provide a data base for sleep studies. Our sample consisted of 174 healthy participants (94 females) aged 20 to 95 years (M = 51.2, SD = 19.6), all of whom provided written informed consent to the respective institute in which their data was collected.

### Recordings

Electroencephalogram (EEG), electrocardiogram (ECG), and airflow (from a respiratory belt) were simultaneously recorded over 2 sessions (2 nights), each around 8 hours long. During recordings, participants were in a supine position with their eyes closed, intending to sleep. For the current study, we extracted data from 3 time periods based on R&K consensus sleep scoring of the EEG data (Anderer et al., 2005). These periods were wakefulness periods immediately prior to sleep, sleep stage 2, and sleep stage 4. We extracted the longest consecutive recording within each period from either night and took the middle 4 minutes of recording within the continuous recording from wakefulness and sleep stage 4, and the middle 10 minutes from sleep stage 2. We excluded participants with less than 2 minutes of maximum consecutive recording in a period, from further analyses in the respective period. In the wakefulness period, we were left with 140 participants (75 females; M_age_ = 52, SD = 19.9, range = [2 90]). In sleep stage 2, we were left with 172 participants (94 females; M_age_ = 50.9, SD = 19.6, range = [20 95]). In sleep stage 4, we were left with 100 participants (59 females; M_age_ = 43.5, SD = 18.9, range = [20 95]).

### Alpha and sleep spindle frequency analysis

We analyzed the EEG data with Matlab code and the Fieldtrip toolbox (Oostenveld et al., 2011). We averaged data from 2 posterior EEG electrodes (O1 and O2). We then computed the derivative of the time-domain signal which results in removing the 1/f component of the frequency-domain signal and enables to better detect peaks nested on the 1/f background. We calculated power spectra based on 1 s segments of data, multiplied with a Hann taper of 1s length, and zero-padded to 10 s. Zero-padding to 10s results in our desired frequency resolution of 0.1 Hz. This approach has been used in studies aiming to detect relatively small differences in peak frequencies (e.g. Haegens et al., 2014). We then applied a Fast-Fourier Transform (FFT) to calculate power between 7 and 14 Hz (AF range) for the wakefulness period and between 11 and 15 Hz for the sleep periods (SF range).

Finally we detected the peak with the highest power using the Matlab function “findpeaks”. We excluded the cases where there were no peaks.

### Heart rate analysis

We used the maximal overlap discrete wavelet transform (MODWT) to enhance the R peaks in the ECG waveform. First, we decomposed the ECG waveform down to level 5 using the Matlab default ‘sym4’ wavelet. Then, we reconstructed a frequency-localized version of the ECG waveform using only the wavelet coefficients at scales 4 and 5, which correspond to the frequency pass-bands known to maximize QRS energy ([11.25 22.5] and [5.625 11.25], respectively; Elgendi et al., 2010). Next we took the squared absolute values of the signal approximation built from the wavelet coefficients and used Matlab’s “findpeaks” to detect the R peaks. This wavelet transform-based method has been shown to efficiently distinguish the QRS complex from high P or T waves, noise, baseline drift, and artifacts (Li et al., 1995). Finally, we counted the number of R peaks in 30s segments of data, averaged the count across segments, and divided by 30 to obtain a value in Hz.

### Breathing frequency analysis

We analyzed the airflow data with Matlab code and the Fieldtrip toolbox. We used the derivative of the time-domain signal to calculate power spectra based on 30 s segments of data, multiplied with a Hann taper of 30s length, and zero-padded to 300 s to achieve a 0.0033 Hz frequency resolution. We then applied a FFT to compute power between a range of 0.05 and 0.5 Hz.

Due to the appearance of multiple breathing rhythms for each participant in the wakefulness data, we narrowed down the frequency range in which to detect peaks in the breathing power spectra. Based on the individual’s heart rate (HR_i_), we calculated a frequency range with a lower limit of ((HR_i_/2)*g)/r and an upper limit of ((HR_i_*2)/g)/r where g is the golden mean (1.618…) and r is the hypothesized HR : BF frequency ratio of 4. For example, for an individual with HR_i_ = 1 Hz, the range would be 0.202 to 0.309 Hz. According to the binary hierarchy brain body oscillation theory (Klimesch, 2018), decoupling between HR and BF would occur at the range’s limits. To include the possibility of obtaining a ratio that would indicate decoupling, we further widened our search-space range by 0.02 Hz at each end. So for an individual with HR_i_ = 1 Hz, we looked for BF peaks within the range of 0.182 to 0.329 Hz.

Finally, we used Matlab’s “findpeaks” to detect the BF peak with the highest power within that range. The sleep data did not exhibit multiple breathing frequencies (“findpeaks” always returned either 0 or 1 peak per participant), so we simply used “findpeaks” to detect the BF peaks within the 0.05 to 0.5 range during sleep stages 2 and 4. We excluded the cases where there were no peaks.

### Statistical analyses

After excluding participants based on the aforementioned criteria, we were left with 109 participants (59 females; M_age_ = 52.5, SD = 20.6, range = [20 95]) during wakefulness. During sleep stage 2, we were left with 124 participants (63 females; M_age_ = 52, SD = 19.7, range = [20 95]). During sleep stage 4, we were left with 73 participants (42 females; M_age_ = 45.2, SD = 19.7, range = [20 95]). We used these data for all statistical analyses.

We had three recording periods (wakefulness, sleep stage 2, and sleep stage 4) and three dependent measures in each (HR, BF, AF/SF) which resulted in 9 sets of frequency ratios to test our 9 hypotheses (see Introduction). To test the empirically obtained ratios against the hypothesized ones, we first used two-sided t-tests, expecting not to reject the null hypotheses (ie to obtain p-values greater than .025). Next, we used the ‘two one-sided tests’ (TOST), an equivalence test whose null hypothesis is defined as an effect large enough to be deemed interesting, specified by an equivalence bound. We set these equivalence bounds for each hypothesized ratio (r) as (r/2)*g for the lower bound, and (r*2)/g for the upper bound. The rationale is that at these limits, a coupling (binary multiple) ratio would become a decoupling (g-related) ratio and vice versa. We expected to reject the null hypotheses of the TOST (both p less than 0.025). Finally, we calculated 95% confidence intervals around the means of each of the 9 distributions using bootstrap sampling from the empirical ratios with 10,000 iterations. We expected that our hypothesized ratios lie within these confidence intervals. We plotted the data using a modified ‘raincloud’ script (Allen et al., 2018).

## Results

In this study, we aimed to test the hypotheses generated by the binary hierarchy brain body oscillation theory (Klimesch 2013, 2018). These hypotheses relate to the ratios (r) obtained from dividing the peak frequencies of brain and body oscillations. We extracted 1 peak frequency value per participant, from each of the EEG, ECG, and airflow (breathing) signals, during periods of wakefulness, sleep stage 2, and sleep stage 4. This totaled 9 frequency ratios, for which we had a hypothesized mean value each. During wakefulness, the expected frequency ratio for AF : HR is r=8, that of AF : BF is r=32, and that of HR : BF is r=4. During both sleep stages, the expected frequency ratio of SF : HR is r=8*g=12.94, that of SF : BF is r=32*g=51.78, and that of HR : BF is r=4. Figures 1-3 summarize the obtained results.

**Figure 1.**
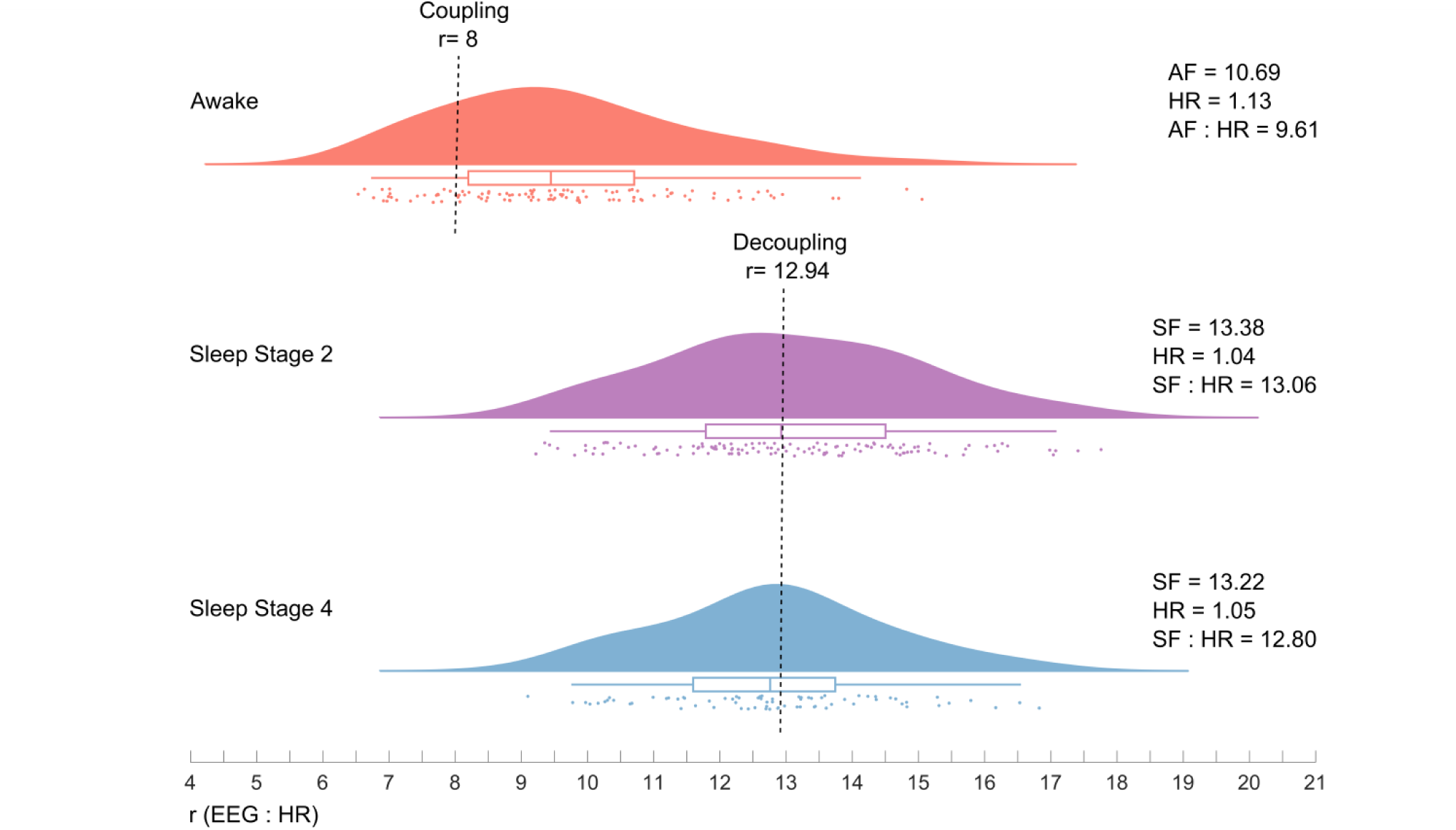
Distribution of peak frequency ratios of individual Alpha Frequency (AF) to Heart Rate (HR) during wakefulness (top panel, red), and Spindle Frequency (SF) to HR during sleep stage 2 (middle panel, purple) and sleep stage 4 (bottom panel, blue). Plots represent individual ratios (jittered dots), probability distribution, and box-plots (median and quartiles). Top: During wakefulness, mean AF : HR was 9.61, significantly different than the predicted coupling ratio of 8. Middle and Bottom: During sleep stage 2, mean SF : HR was 13.06, not significantly different than the predicted decoupling ratio of 12.94. Bottom: During sleep stage 4, mean SF : HR was 12.8, not significantly different than the predicted decoupling ratio of 12.94.

For the AF : HR and SF : HR ratios (Figure 1), we did not find the predicted ratio of 8 during wakefulness (M=9.61, SD=1.85, CI = [9.26 - 9.99], t(108) = 6.46, p<.001, p_TOST_=.78), but we did find the predicted ratio of 12.94 during both sleep stage 2 (M=13.06, SD=1.93, CI = [12.72 – 13.37], t(123)=0.5, p=.97, p_TOST_<.001) and sleep stage 4 (M=12.8, SD=1.69, CI = [12.43 – 13.24], t(72)=-0.52, p=.97, p_TOST_<.001).

For the AF : BF and SF : BF ratios (Figure 2), we did not find the predicted ratio of 32 during wakefulness (M=38.07, SD=7.85, CI = [36.63 – 39.64], t(108)=5.72, p<.001, p_TOST_=.64), but we did find the predicted ratio of 51.78 during both sleep stage 2 (M=52.31, SD=7.72, CI = [50.96 – 53.59], t(123)=0.59, p=.98, p_TOST_<.001) and sleep stage 4 (M=51.28, SD=7.47, CI = [49.45 – 53.38], t(72)=-0.4, p=.99, p_TOST_<.001).

**Figure 2.**
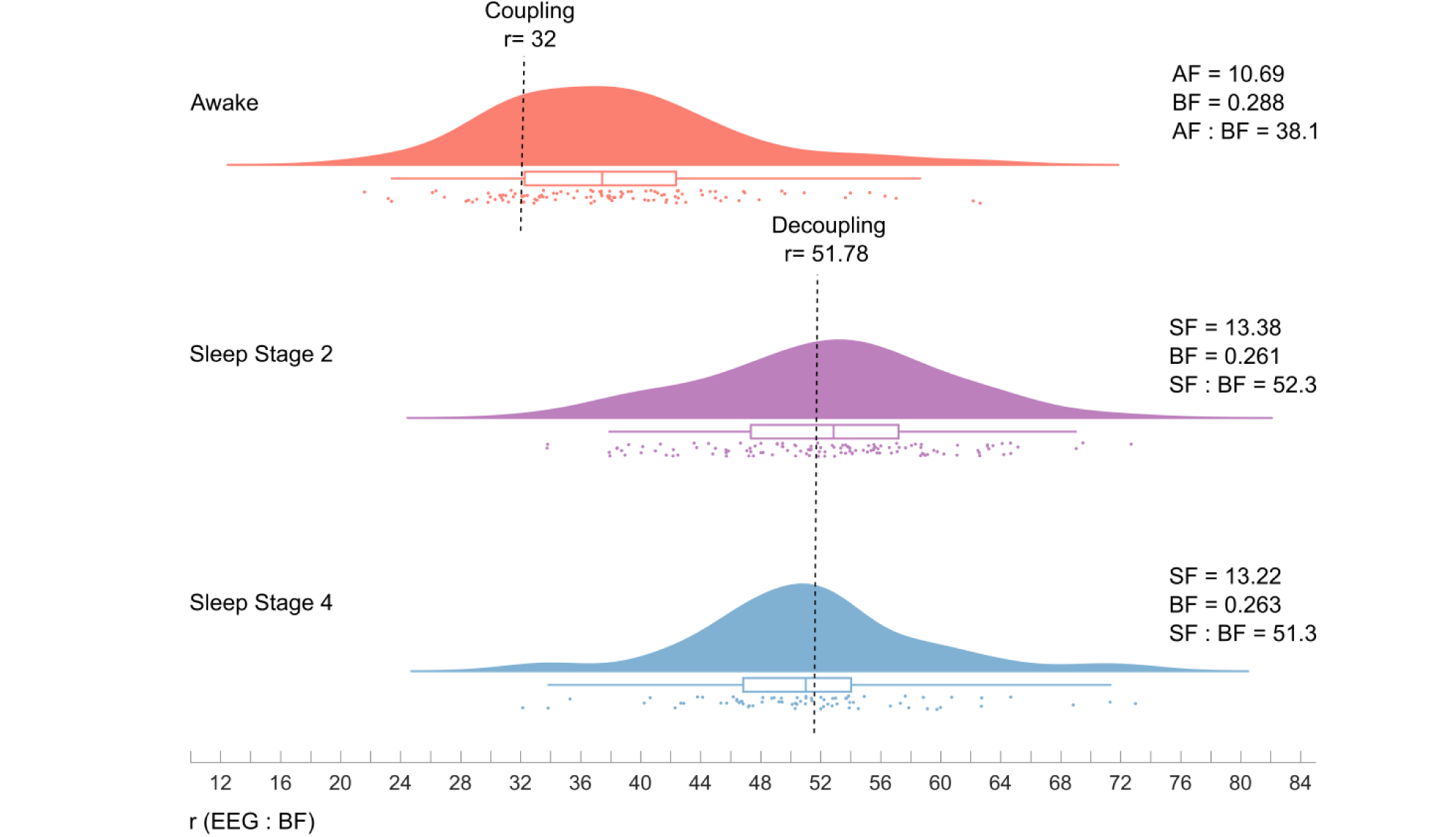
Distribution of peak frequency ratios of individual Alpha Frequency (AF) to Breathing Frequency (BF) during wakefulness (top panel, red), and Spindle Frequency (SF) to BF during sleep stage 2 (middle panel, purple) and sleep stage 4 (bottom panel, blue). Plots represent individual ratios (jittered dots), their probability distribution, and box-plots (median and quartiles). Top: During wakefulness, mean AF : BF was 38.1, significantly different than the predicted coupling ratio of 32. Middle and Bottom: During sleep stage 2, mean SF : BF was 52.3, not significantly different than the predicted decoupling ratio of 51.78. Bottom: During sleep stage 4, mean SF : BF was 51.3, not significantly different than the predicted decoupling ratio of 51.78.

For the HR : BF ratios (Figure 3), we found the predicted ratio of 4 during all stages of wakefulness (M=3.98, SD=0.46, CI = [3.89 - 4.06], t(108)=-0.34, p=.99, p_TOST_<.001), sleep stage 2 (M=4.03, SD=0.45, CI = [3.94 – 4.12], t(123)=0.53, p=.98, p_TOST_<.001), and sleep stage 4 (M=4.02, SD=0.45, CI = [3.91– 4.14], t(72)=-0.48, p=.98, p_TOST_<.001).

**Figure 3.**
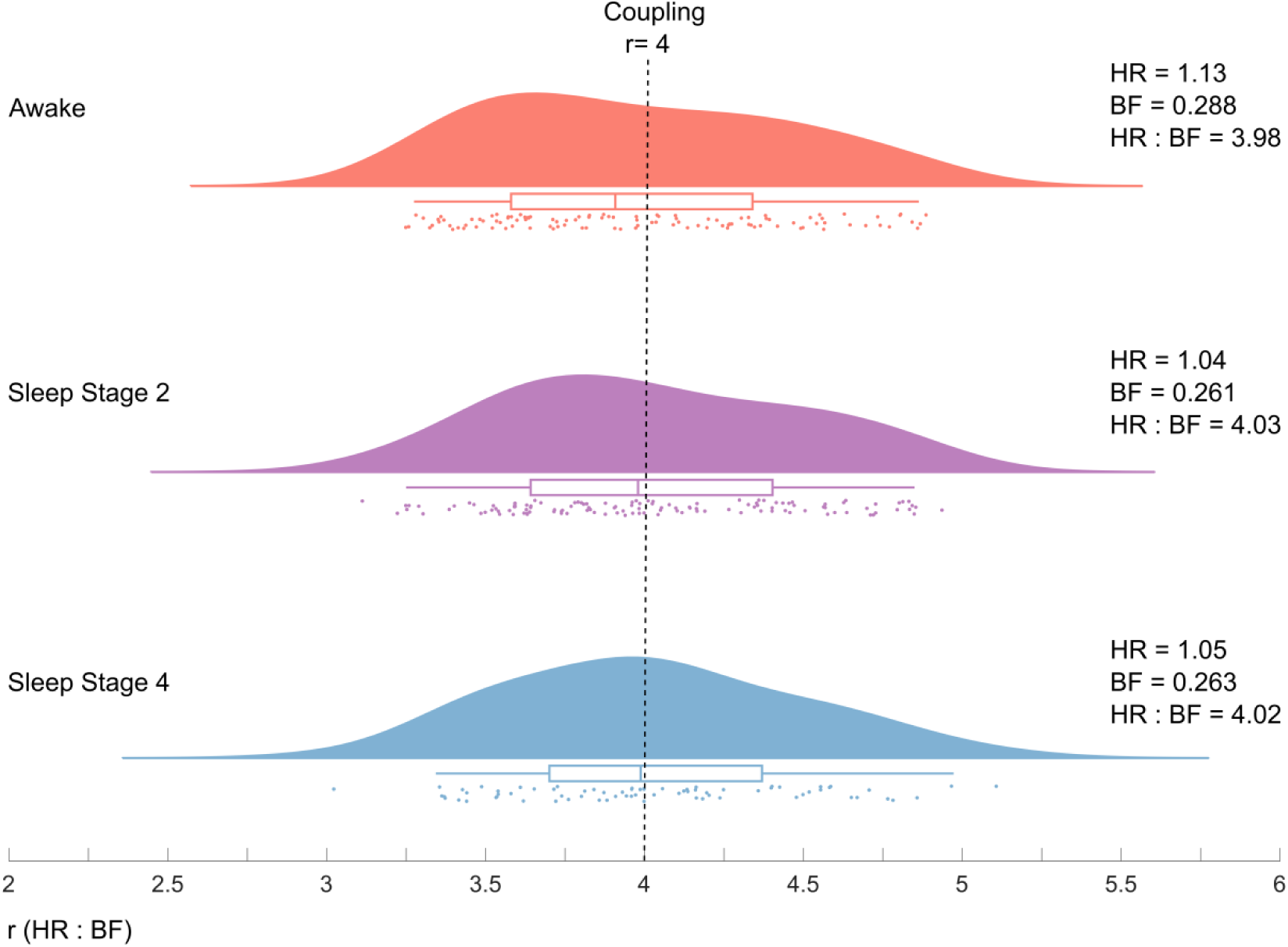
Distribution of peak frequency ratios of individual Heart Rate (HR) to Breathing Frequency (BF) during wakefulness (top panel, red), sleep stage 2 (middle panel, purple) and sleep stage 4 (bottom panel, blue). Plots represent individual ratios (jittered dots), their probability distribution, and box-plots (median and quartiles). Top: During wakefulness, mean HR : BF was 3.98, not significantly different than the predicted coupling ratio of 4. Middle and Bottom: During sleep stage 2, mean HR : BF was 4.03, not significantly different than the predicted coupling ratio of 4. Bottom: During sleep stage 4, mean HR : BF was 4.02, not significantly different than the predicted coupling ratio of 4.

In sum, we found supportive evidence for 7 out of our 9 hypotheses.

## Discussion

The binary hierarchy brain body oscillation theory (Klimesch, 2013, 2018) refers to two groups of predictions. (i) During alert wakefulness binary multiple frequency ratios are expected between brain and body oscillations (AF: HR, AF : BF) and body oscillations (HR : BF). (ii) During sleep, irrational ratios are expected between brain and body oscillations (SF: HR, SF : BF), but a binary multiple ratio for body oscillations (HR : BF). The findings of our study support all predictions for sleep but only one prediction (regarding the expected ratio of 1: 4 for HR : BF) for wakefulness. The observed ratios between AF : HR and AF : BF (of 9.61 and 38.07) deviated significantly from the predicted ratios (of 8 and 32). We suspect that this negative finding is due to the nature of the sleep study from which the data were taken. Subjects were recorded while they were awake but with their eyes closed, intending to fall asleep. The wakefulness epochs of data were extracted from the period immediately preceding falling asleep. While people were physiologically awake, it is very likely that there was no active cognitive processing or movement during these periods, and that HR and BF had already dropped below the average values one would expect during active wakefulness. Inspection of mean HR (see Figures, right side) supports this interpretation. During wakefulness, HR (equaling 1.13 Hz) is only slightly faster than in stage 2 and stage 4 sleep, where HR drops to 1.04 Hz. Textbooks (e.g. Asharaya et al., 2007) and studies measuring mean HR in large samples (e.g. Shaffer et al. 2014) show higher values around 1.25 Hz during rest. It is possible that the observed ratios of 9.61 and 38.07 (for AF : HR and AF : BF) already reflect a shift towards the predicted irrational ratios of 12.94 and 51.

We found that sleep spindle frequency is irrationally related to body oscillations (HR and BF), whereas the harmonic ratio between the two body oscillations remained constant throughout wakefulness and sleep. We argue that this finding reflects a situation where coupling between brain and body oscillations is suppressed, possibly reflecting decreased body awareness.

With respect to BF it is important to note that we observed multiple peaks in the spectra of the breathing data during wakefulness and selected only those peaks which fell within the frequency range predicted by HR. This means that during rest, multiple breathing frequencies were present, from which one frequency exhibits the predicted ratio of 4. During sleep, however, all spectra exhibited one peak only. This suggests that coupling between HR and BF is stronger during sleep as compared to wakefulness.

Multiple breathing frequencies were also reported in research carried out by Perlitz, Lambertz and colleagues. These authors report evidence for a ‘0.15 Hz’ rhythm (e.g., Perlitz et al., 2004, Lambertz & Langhorst, 1998; Lambertz et al., 2000) which can be observed primarily during periods of relaxation. Most importantly, Perlitz et al., (2004) have found that BF entrains to this 0.15 Hz rhythm at integer frequency ratios of 1:1, 2:1 or 1:2. Accordingly, BF exhibits a 1:1 or doubling/halving relationship relative to the 0.15 Hz rhythm with dominant frequencies at around 0.15 Hz, 0.30 Hz and 0.075 Hz (cf. Tab.1 in Perlitz et al. 2004). These frequencies reflect binary ratios relative to HR (HR/4 = 0.313; HR/8 = 0.156; HR/16 = 0.078), and thus provide additional evidence for a binary hierarchy of oscillations, as suggested by Klimesch (2018).

Taken together, the findings suggest that frequency ratios are sensitive measures, reflecting different frequency architectures that enhance or suppress coupling. For an empirical evaluation it is critical to define situations where one can predict coupling or de-coupling. For the active brain and body, coupling and de-coupling may occur transiently, for short time periods, and for selected oscillations only. Thus, it is methodologically difficult to detect frequency coupling or de-coupling of brain-body oscillations during active task performance. However, sleep - and deep sleep in particular – provides an almost ideal situation for the investigation of our hypotheses, because of the complete lack of external task demands and because brain and body oscillations are modulated endogenously over longer time periods. We conclude that the observed frequency ratios between SF and HR, and SF and BF, provide good evidence for decoupling between brain and body oscillations during sleep.

## Author contributions

E.R. and W.K. conceived the project and wrote the manuscript. E.R. analyzed the data. G.D. provided the data and consulted on writing the manuscript. W.G. consulted on statistical analyses.

## Acknowledgements

This work was supported by the FWF (Austrian Science Fund) grant: Imaging the Mind: Connectivity and Higher Cognitive Function (W 1233-G17).

## Data and code availability

The raw data used to extract the variables analyzed in this study are not publicly available and were used under license for this study. Processed variables from the data as well as computer code used are available upon request to the corresponding author E.R.

